# HNF4A is required to specify glucocorticoid action in the liver

**DOI:** 10.1101/2021.04.10.438998

**Authors:** A. Louise Hunter, Toryn M. Poolman, Donghwan Kim, Frank J. Gonzalez, David A. Bechtold, Andrew S. I. Loudon, Mudassar Iqbal, David W. Ray

## Abstract

The glucocorticoid receptor (GR) is a nuclear hormone receptor critical to the regulation of energy metabolism and the inflammatory response. The actions of GR are highly dependent on cell type and environmental context. Here, we demonstrate the necessity for liver lineage-determining factor hepatocyte nuclear factor 4A (HNF4A) in defining liver-specificity of GR action. In normal mouse liver, the HNF4 motif lies adjacent to the glucocorticoid response element (GRE) at GR binding sites found within regions of open chromatin. In the absence of HNF4A, the liver GR cistrome is remodelled, with both loss and gain of GR recruitment evident. Lost sites are characterised by HNF4 motifs and weak GRE motifs. Gained sites are characterised by strong GRE motifs, and typically show GR recruitment in non-liver tissues. The functional importance of these HNF4A-regulated GR sites is further demonstrated by evidence of an altered transcriptional response to glucocorticoid treatment in the *Hnf4a*-null liver.

## Introduction

NR3C1, the glucocorticoid receptor (GR), is an almost ubiquitously expressed nuclear receptor. Whilst there is evidence for rapid, non-genomic actions of glucocorticoids (1, 2), the chief outcomes of GR activation occur through its nuclear activity. GR binds the genome through the glucocorticoid response element (GRE) motif, which comprises two palindromic hexamers separated by a 3bp spacer (AGAA-CANNNTGTTCT). Formation of higher order GR structures (dimerisation, tetramerisation) is necessary for GR-mediated gene regulation (3).

Upon ligand binding, GR is predominantly directed to sites on the genome where chromatin is already accessible (4), and this is dependent on cell type, with priming by C/EBPB (CCAAT-enhancer binding protein beta) particularly important in the liver (5). GR can also demonstrate pioneer function at sites of inaccessible chromatin (6). Whilst it is less clear what the determinants of binding are here, similarity of the GR-bound DNA sequence to the canonical GRE (’motif strength’) may play a role, with nucleosome-deplete sites demonstrating more degenerate GRE motifs (6). Other studies have shown that active histone marks and presence of pioneer factors also play a role in dictating GR binding (7). Following GR binding, gene activation - involving recruitment of coactivators and chromatin remodellers - occurs at sites of pre-established enhancer-promoter interactions, with the presence of GR increasing the frequency of productive interactions (8). In contrast, the mechanism by which GR downregulates gene expression remains an area of considerable debate, with evidence for protein-protein tethering, indirect mechanisms of action, and GR binding to negative or cryptic response elements presented (9–14).

Surprisingly for a transcription factor which is so widely expressed, GR action is remarkably context-specific. GR activity can be influenced both by metabolic and immune state (15–17). GR action is also highly tissue-specific, with the GR cistrome showing limited overlap between different cell types (4). This is a property which is far from unique to GR, and has been well-illustrated for other transcription factors from multiple classes, including the oestrogen receptor (18), and the core clock protein BMAL1 (19). *In vitro* studies suggest that GR cell-specificity is conferred by differential chromatin accessibility at distal enhancer sites, with GR binding in proximal promoter regions regulating genes which are ubiquitously GC-responsive (20). In this *in vivo* study, we show the dominance of hepatocyte nuclear factor 4A (HNF4A), itself a nuclear receptor, in determining GR binding in mouse liver. We find the HNF4 motif to underlie sites of GR binding, and, in *Hnf4a*-null liver, demonstrate loss of GR binding at HNF4-marked sites and emergence of new, non-liver-specific GR binding events at sites characterised by strong GRE motifs. This remodelling of the GR cistrome is further demonstrated to be of functional importance, in shaping an altered transcriptomic response to glucocorticoids in the absence of HNF4A.

## Results

### HNF4 motifs mark liver GR binding sites

We first mapped the hepatic GR cistrome by performing GR ChIP-seq on mouse liver collected one hour after intraperitoneal injection of dexamethasone (DEX) at Zeitgeber Time 6 (mice housed in 12hr:12hr light-dark cycles, ZT0 = lights on) (Fig.1A). As expected, DEX treatment resulted in substantial GR recruitment to the genome, with 20,064 peaks called over input (q<0.01) in DEX-treated tissue.

**Fig. 1.**
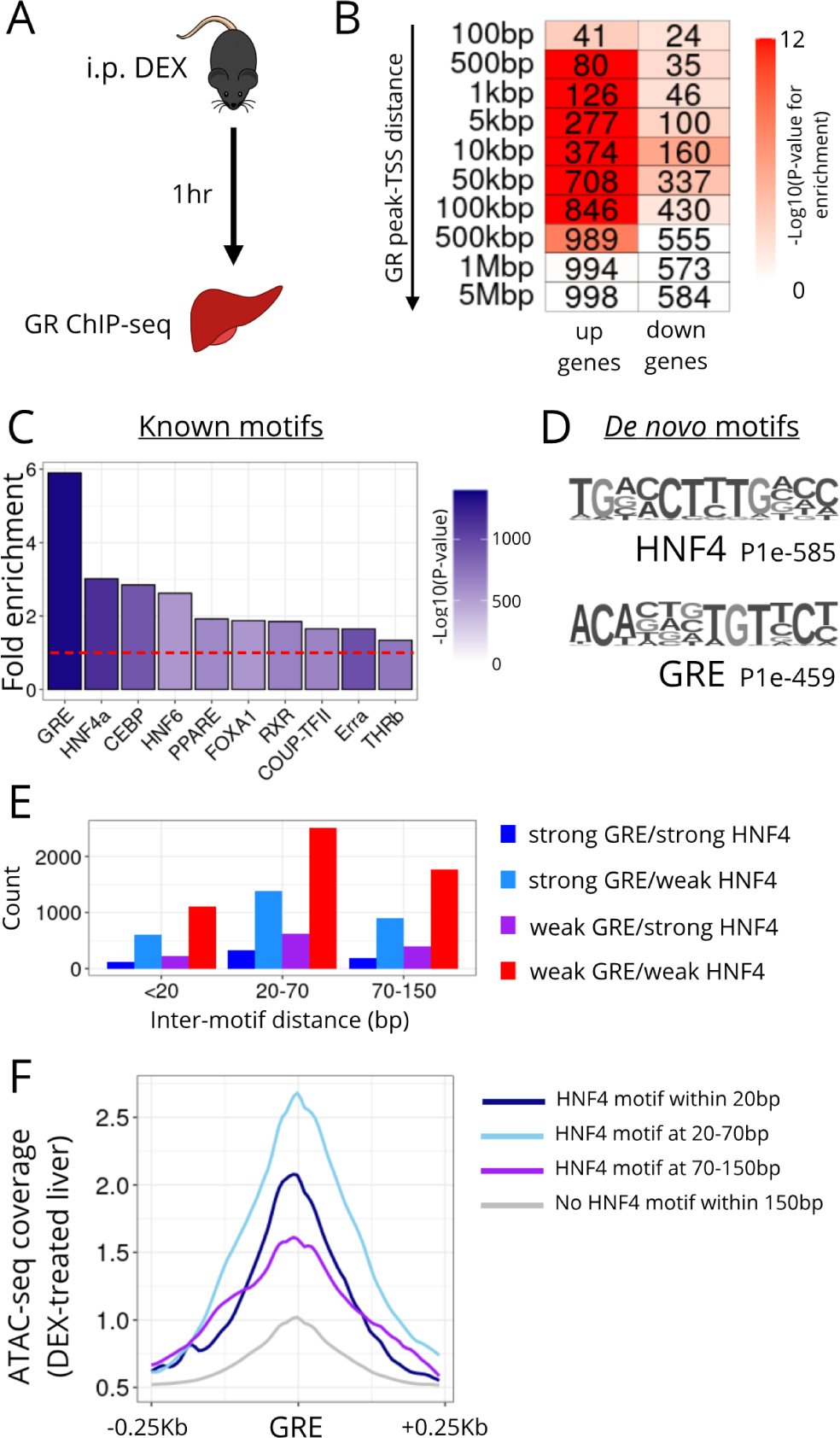
Liver GR binding sites are marked by GRE and HNF4A motifs. **A.** Liver GR ChIP-seq was performed one hour after acute dexamethasone (DEX) treatment. **B.** Heatmap showing enrichment (hypergeometric test) of the transcription start sites (TSSs) of genes up or downregulated by glucocorticoid treatment at increasing distances from GR ChIP-seq peaks. Shading of each cell indicates −log10(P-value) for enrichment (over all genes in the genome), number indicates number of genes in each cluster at that distance. **C.** Fold enrichment, in GR ChIP-seq peaks, of known motifs. Red dotted line at y=1. **D.** The two motifs detected most strongly (lowest P values) *de novo* in GR peaks. **E.** Barchart of inter-motif distances for GRE and HNF4 motifs detected within GR ChIP-seq peaks. **F.** ATAC-seq coverage score (mean coverage from 2 biological replicates), in DEX-treated liver, around canonical GRE motifs with or without a HNF4 motif within specified distances.

We have previously shown that the same model of gluco-corticoid treatment has a large effect on the liver transcriptome, with the expression of 1,709 genes being significantly altered (the majority upregulated) (21). We now observed pronounced enrichment of glucocorticoid-upregulated genes in relation to GR binding sites (Fig.1B; hypergeometric test (22)). Enrichment was seen at distances of 500bp-500kbp between transcription start sites (TSS) of DEX-activated genes and GR ChIP-seq peaks (Fig.1B), implying both proximal and distal regulation. The noticeably weaker enrichment of glucocorticoid-downregulated genes supports the notion, reported by others (11, 23), that gene downregulation occurs by indirect means; however, we cannot exclude mechanisms such as tethering based on these data.

We proceeded to motif discovery analysis, and found the canonical GRE to be the most highly-enriched known motif within GR binding sites, followed by the HNF4 motif (Fig.1C, see also [Mendeley Data](https://data.mendeley.com/datasets/k8d386ndz6/2)). A motif most closely resembling the HNF4 motif had the lowest P-value on *de novo* motif analysis (Fig.1D), and was detected in 25.64% of GR peaks. As reported for other GR ChIP-seq studies (24, 25), other motifs detected (at lower significance) included CEBP, PPAR and HNF6 motifs. A similar pattern of motif enrichment was seen in GR ChIP-seq peaks called in vehicle-treated liver (5,831 peaks called over input), with the known GRE and HNF4 motifs being the most strongly enriched, and a HNF4-like motif detected *de novo* in 29.29% of peaks (Fig.S1A–B). Interestingly, we still observed strong enrichment of DEX-upregulated genes at distances of 5-500kbp from GR peaks in VEH liver (Fig.S1C), implying some pre-existing GR binding in association with glucocorticoid-responsive genes.

We were interested to know how closely GRE and HNF4 motifs were situated at GR binding sites, as the distance between transcription factor (TF) motifs contributes to the likelihood of TFs co-occupying regulatory elements (26), with a random distribution of co-occupancy events observed at inter-motif distances >70bp, and co-occupancy being most likely at distances <20bp. High levels of co-occupancy could suggest that binding is physically co-operative (26). Under GR peaks where we observed co-occurrence of HNF4 and GRE motifs, we saw a spread of inter-motif distances (Fig.1E), with the majority in the range of 20-70bp, irrespective of whether motif calling was performed with high stringency settings (“strong” GRE/HNF4), or permitted some degeneracy (“weak“). Thus, these data favour co-occupancy of GR and HNF4A at regulatory elements, but do not necessarily support physical co-operation between the two nuclear receptors.

To explore the importance of HNF4A via an independent approach, we performed ATAC-seq (assay for transposase-accessible chromatin) on the same DEX-treated samples of liver. We used HOMER to map the positions of all canonical GREs in the mouse genome (>137,000 locations). At GREs with a nearby HNF4 motif (within 20bp, n=3,281; at 20-70bp, n=2,986; at 70-150bp, n=4,872), we observed stronger mean ATAC signal than at GREs without a nearby HNF4 motif (n>125,000) (Fig.1F), with signal strongest at 20-70bp inter-motif distances. Therefore, in liver, HNF4A plays a role in determining chromatin accessibility at GR binding sites.

Taken together, the above data show that HNF4A is critical in determining the liver GR cistrome. Co-located GRE and HNF4 motifs mark regulatory elements where GR acts to upregulate gene expression in the *cis* domain. This is in line with existing theories that GR binds the genome at pre-programmed, DNase-sensitive sites (4, 5, 27), with glucocorticoid treatment augmenting GR action at these sites, and increasing the frequency of pre-established enhancer-promoter interactions (8).

### HNF4A loss remodels the GR cistrome

We thus hypoth-esised that removing HNF4A would impact upon patterns of GR binding. To test this, we performed GR ChIP-seq on livers from 6-8 week-old *Hnf4a^fl/fl^Alb^Cre^* mi ce (28) treated acutely with DEX, again at ZT6. *Hnf4a^fl/fl^Alb^Cre+/−^* mice are viable, but show hepatomegaly and hepatosteatosis from 6 weeks of age, and increased mortality from 8 weeks of age (28). Nonetheless, they present a useful model to study how HNF4A regulates transcriptional activity *in vivo*. We performed a differential binding analysis (29, 30) to detect sites where GR binding was statistically different (FDR<0.05) between *Hnf4afl/flAlbCre-/-* (Cre-) and *Hn f4a^fl/fl^Alb^Cre+/−^* (Cre+) livers. We employed an internal spike-in normalisation strategy with *D.melanogaster* chromatin (31) to control for technical variation, and so increase confidence that results represented true genotype effects.

This approach detected 4,924 sites where GR binding was lost in Cre+ animals compared to Cre-, and 989 sites where GR binding was gained. Loss of HNF4A therefore led to substantial remodelling of the liver GR cistrome (Fig.2A,B). Lost and gained sites chiefly annotated to intergenic and in-tronic regions of the genome, suggesting that remodelling was affecting distal regulatory sites rather than proximal promoter regions (Fig.S2A). In keeping with previous work (20), this supports the notion that tissue-specificity of GR action may be conferred by distal enhancers.

**Fig. 2.**
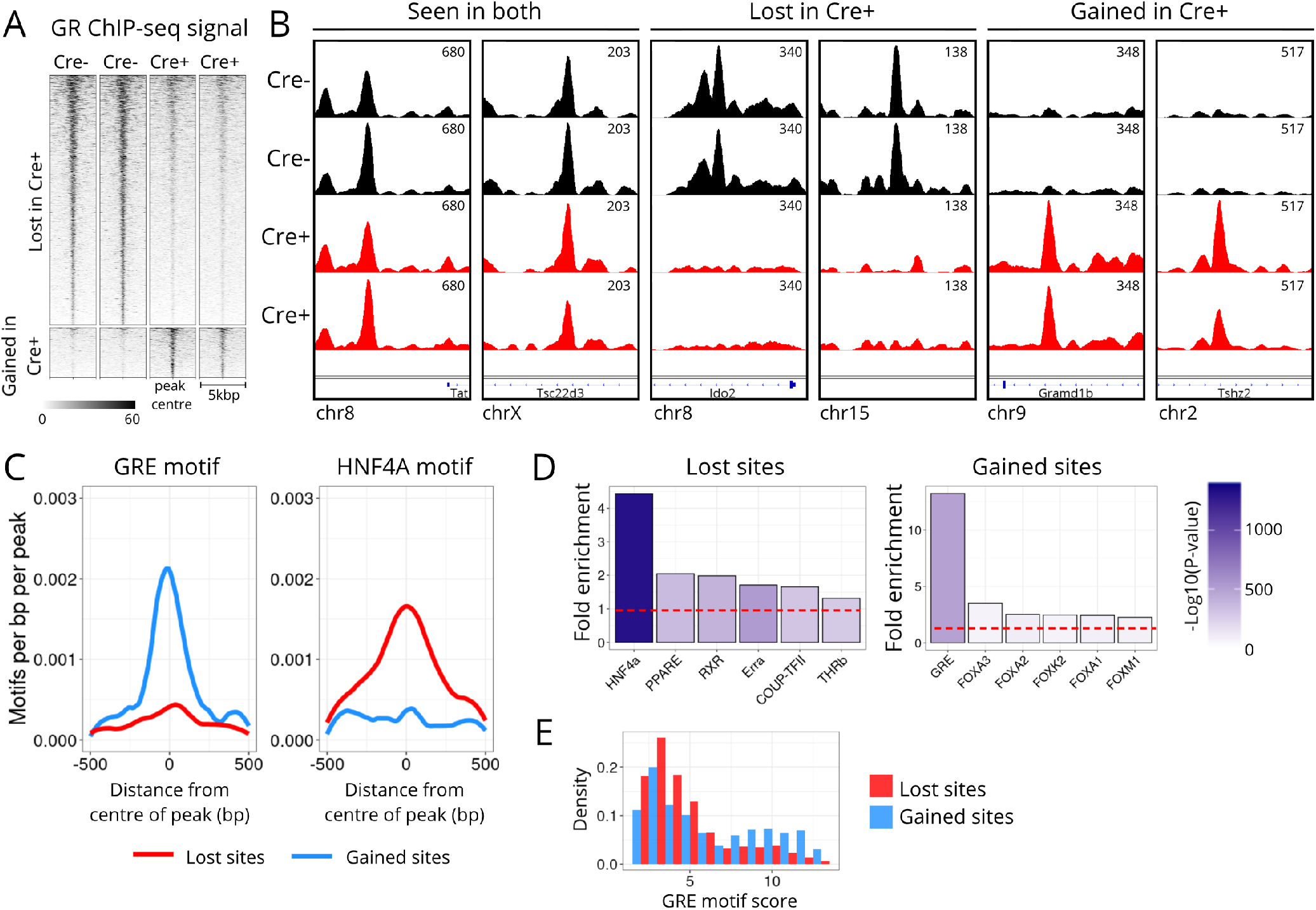
The liver GR cistrome is remodelled in the absence of HNF4A. **A.** GR ChIP-seq signal at *csaw*-detected DB sites in *Hnf4a^fl/fl^Alb^Cre^* Cre- and Cre+ mouse liver (signal shown +/− 2.5kbp from centre of each GR site). **B.** GR ChIP-seq signal tracks of exemplar DB sites. Y axis is uniform within each panel. **C.** Abundance of the GRE and HNF4A motifs within lost (red) and gained (blue) GR sites. **D.** Fold enrichment, in lost and gained GR sites, of the 6 most highly enriched known motifs. Red dotted line at y=1. **E.** Density histogram of GRE motif scores (measure of motif strength) in lost (red) and gained (blue) GR sites.

To examine the distinctions between lost and gained GR sites in more detail, we first quantified abundance of specific motifs. We observed that GR sites lost in Cre+ liver showed low abundance of the canonical GRE and high abundance of the HNF4 motif, whilst GR sites gained showed high abundance of the canonical GRE and low HNF4 motif abundance (Fig.2C). These findings were recapitulated by motif discovery analysis (Fig.2D, see also [Mendeley Data](https://data.mendeley.com/datasets/k8d386ndz6/2)), with the enrichment of the HNF4 motif in lost GR sites supporting the specificity of the effect. Comparison of our data with recently published HNF4A liver cistromes demonstrated overlap of lost GR sites with HNF4A binding sites (Fig.S2B), suggesting that the HNF4A protein, in addition to the motif, can normally be found at these sites. Unsurprisingly, we saw almost no overlap of gained GR sites with the HNF4A cistrome. On comparing the strength of GRE motifs in lost and gained sites (scored by similarity to the canonical GRE), we found lost GR sites to be predominantly characterised by weak GREs, whilst strong GREs characterised a population of gained sites (Fig.2E). Within lost sites, co-occurrence of GRE and HNF4 motifs was most numerous at inter-motif distances of 20-70bp (Fig.S2C), as we observed for the wider GR cistrome.

Therefore, in the absence of HNF4A, GR is no longer recruited to sites marked by the HNF4 motif (and HNF4A binding), and a weak GRE motif. Intriguingly, GR binding emerges at sites where it is not normally recruited, where strong GRE motifs are present (and where HNF4A is not found). This marked divergence between lost and gained sites points to this being a specific consequence of HNF4A loss, and not a downstream effect of the abnormal hepatic physiology of *Hnf4a^fl/fl^Alb^Cre+/−^* livers.

### HNF4A-dependent GR sites demonstrate distinct patterns of chromatin accessibility and tissue GR recruitment

Next, we sought to understand the normal chromatin state of GR sites lost or gained in the absence of HNF4A. We took advantage of published datasets for DNase hypersensitivity (32) and histone mark ChIP-seq (33, 34) in naive mouse liver (ie. the chromatin landscape which the GR encounters upon dexamethasone treatment) to quantify read coverage at lost and gained sites. We found that GR is lost at sites where, in liver, chromatin is normally DNase-sensitive, and where higher levels of the histone marks H3K27ac and H3K4me1 - associated with active/poised enhancers - are found. GR binding is gained at sites where chromatin is normally less DNase-sensitive, and where the H3K27me3 mark (associated with inactive regions of heterochromatin) is stronger (Fig.3A).

**Fig. 3.**
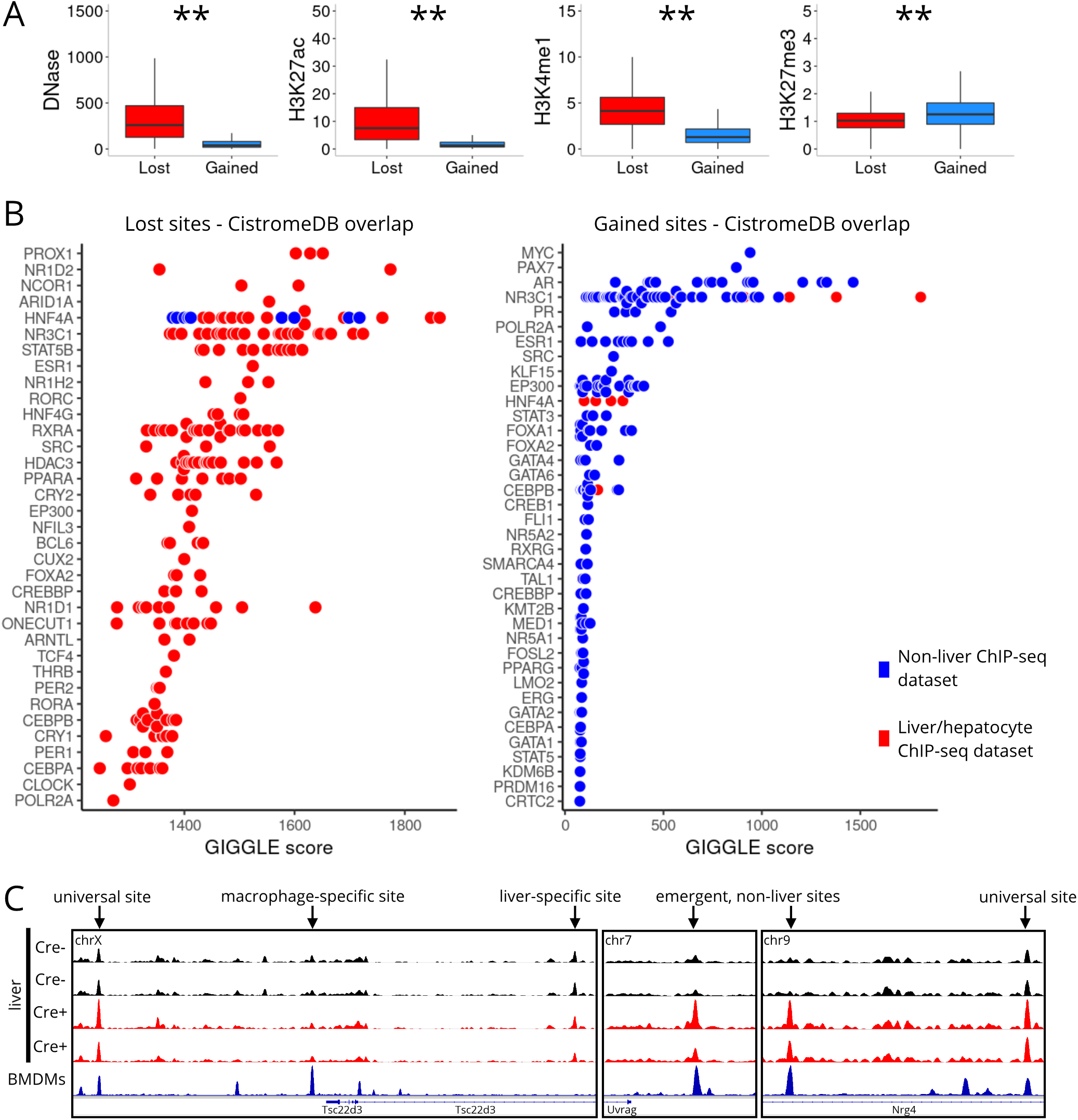
Lost and gained GR sites diverge by chromatin state and tissue-specificity. **A.** Box-and-whisker plots showing read coverage of lost and gained GR sites of signal from DNase-seq and ChIP-seq of histone marks H3K27ac, H3K4me1, H3K27me3. **P<0.01, Wilcoxon tests. Central line at median, box limits at 25th and 75th percentiles, whiskers extend 1.5x interquartile range from box limits. **B.** Overlap of lost and gained GR sites with published transcription factor cistrome data (top 1k peaks in each dataset), as determined and scored by GIGGLE. Datasets from non-liver tissues/cells plotted in blue, datasets from liver/hepatocytes plotted in red. **C.** Exemplar tracks showing GR ChIP-seq signal around the *Tsc22d3* (*Gilz*), *Uvrag*, and *Nrg4* loci in Cre- and Cre+ liver, and in bone marrow-derived macrophages (11). Universal, macrophage-specific and liver-specific GR sites highlighted by arrows. Y axis is uniform within each panel.

Given that HNF4A is a lineage-determining transcription factor, we hypothesised that loss of HNF4A results in a GR cistrome less specific to the liver. We performed an unbiased comparison of lost and gained sites with transcription factor cistromes, using the GIGGLE tool (35). We found lost sites to be normally bound by not only HNF4A itself, but by multiple other factors with important metabolic roles, with almost all of these cistromes being derived from liver/hepatocyte experiments (Fig.3B, see also [Mendeley Data](https://data.mendeley.com/datasets/k8d386ndz6/2)). In marked contrast, gained sites were most numerously bound by GR (NR3C1) and other NR3 family members, but almost exclusively in non-liver tissues. Importantly, these tissues were diverse, and not simply non-hepatocyte cell types (e.g. inflammatory cells) which might be found within liver tissue. Interestingly, those liver GR cistromes which did show overlap with gained sites (Fig.3B) were from a study where over-expression of a dominant negative form of C/EBP was used to disrupt the liver GR cistrome (5)

Thus, these findings suggest that HNF4A is necessary for GR binding at liver-specific sites, by means of maintaining open chromatin. The GREs at these HNF4A-dependent sites show degeneracy from the canonical AGAACANNNTGTTCT motif, but chromatin state is favourable towards GR binding, being marked by active histone modifications. It may be that other important regulators of liver energy metabolism are also recruited to these regions. By contrast, in the absence of HNF4A, GR is recruited to additional sites where chromatin is not normally accessible in liver, but where a strong GRE motif is found, and where GR is capable of binding in other tissues (Fig.3C).

### HNF4A loss remodels the glucocorticoid-responsive transcriptome

To examine the functional importance of GR cistrome remodelling, we then performed liver RNA-seq to quantify gene expression in dexamethasone- and vehicle-treated *Hnf4a^fl/fl^Alb^Cre^* mice (n=3-4/group). Simply by studying differential gene expression in vehicle-treated Cre+ and Cre- mice, we saw a profound effect of HNF4A on the liver transcriptome (Fig.4A), with the expression of >7,000 genes being different between genotypes. Genes with diminished expression in Cre+ mice were characterised by pathways of lipid, amino acid and oxidative metabolism, whilst genes with increased expression in Cre+ mice were associated with Rho GTPase (intracellular actin dynamics) and cell cycle pathways (Fig.4B).

**Fig. 4.**
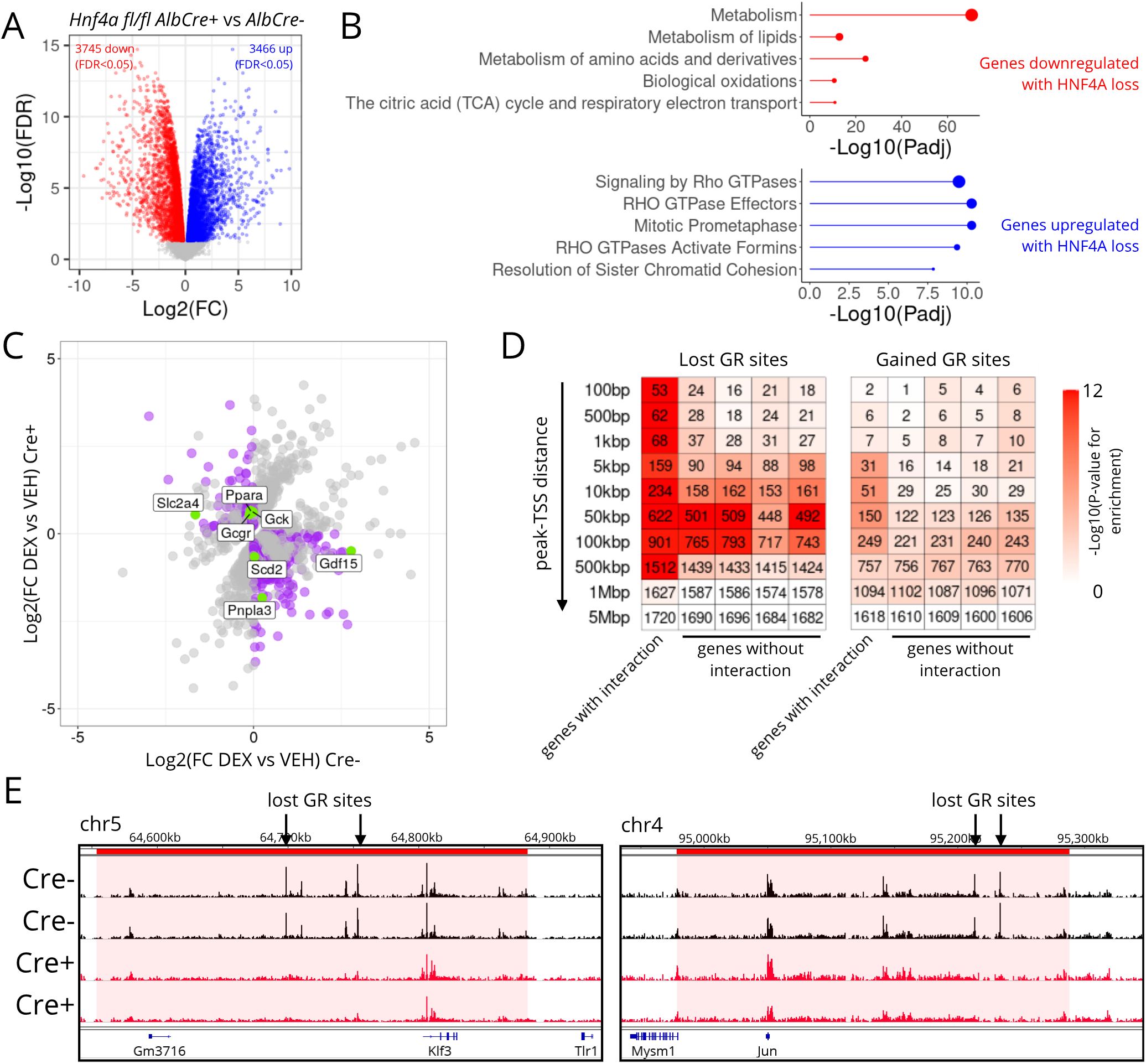
Lost and gained GR sites associate with gene showing altered glucocorticoid response. **A.** Liver RNA-seq in vehicle-treated *Hnf4a^fl/fl^Alb^Cre^* mice, Cre+ vs Cre- samples. Significantly downregulated genes (FDR<0.05) in red, significantly upregulated genes in blue. **B.** Top Reactome pathways of genes downregulated (red) and upregulated (blue) by HNF4A loss. Point size is proportional to number of genes in that pathway. **C.** Effect of DEX treatment in Cre- and Cre+ mice. Genes where stageR detects a significant treatment x genotype interaction shown in grey (n=1,908). Those where direction of change is different between genotypes, and where effect of treatment is significant, highlighted in purple (n=633). These include notable metabolic regulators and enzymes, highlighted in green. **D.** Enrichment of gene clusters at increasing distances from lost or gained GR sites (hypergeometric test (22)). First cluster comprises those 1,908 genes where stageR detects a treatment-genotype interaction, other clusters comprise random samples of equivalent size (repeated x4) of DEX-responsive genes where no treatment-genotype interaction is detected. Shading of each heatmap cell corresponds to −Log10(P-value) for enrichment, number indicates number of genes in each cluster at that distance. **E.** Exemplar tracks showing GR ChIP-seq signal around the *Klf3* and *Jun* loci in Cre- and Cre+ liver. Lost GR sites indicated. Red shading shows the dimensions of the encompassing sub-topologically associating domain (subTAD). subTAD coordinates from (36). Y axis is uniform within each panel.

Importantly, loss of HNF4A also altered the response to acute glucocorticoid treatment. By comparing gene expression in Cre+ and Cre- mice treated with dexamethasone or vehicle, using R Bioconductor package stageR (37), specifically designed to control false discovery rate at the gene level, we found 1,908 genes where a significant genotype-treatment interaction was detected (adjusted P-value <0.05). Of these 1,908 genes, 633 showed a marked difference in response to DEX treatment between the two genotypes (Fig.4C). Of note, these included important metabolic genes (e.g. *Ppara*, *Gdf15*, *Gck*), suggesting that HNF4A exerts a major impact on the liver metabolic response to glucocorticoids. *Gdf15* expression, for example, is normally upregulated by glucocorticoid, an effect which is lost in the *Hnf4a^fl/fl^Alb^Cre+/−^* mice.

To determine whether these changes in transcriptomic response might directly relate to the remodelling of the GR cistrome, we looked for significant enrichment of genes of interest in the locale of lost and gained GR sites. Specifically, we looked at the 1,908 genes where stageR detected a significant treatment-genotype interaction, and asked whether these were over-represented in proximity to GR sites (Fig.4D). As control clusters, we took DEX-responsive genes where stageR did not detect such an interaction (n=3,797) and randomly sampled 1,908 genes from this group, repeating this random sampling four times.

At GR sites lost in the absence of HNF4A (n=4,924), at distances between 100bp-500kbp, we saw strong enrichment of genes with a treatment-genotype interaction (e.g. *Klf3* and *Jun*, Fig.4D,E). Interestingly, strong enrichment of genes without an interaction was also apparent, but at distances of 50-100kbp from lost GR sites (Fig.4D). It may be that a proportion of these sites are distal enhancers which make a redundant contribution to the regulation of the genes in question, or a contribution that is sufficiently small not to be detected in our RNA-seq analysis. Other, HNF4A-independent regulatory elements may exert more dominant control over these genes. For GR binding sites newly emergent in Cre+ liver (gained sites, n=989), we saw distinct enrichment of small numbers of genes with a treatment-genotype interaction at distances of 5-100kbp, with this pattern not observed for genes without an interaction (Fig.4D). These results are consistent with the idea that loss and gain of GR binding in *Hnf4a*-null liver contributes to the observed alteration in the transcriptional response to glucocorticoid, and that the remodelling of the GR cistrome is of functional importance.

### HNF6 has limited influence on liver GR action

We were also interested to examine the influence of a hepatic lineage-determining factor from another family. We have found HNF4A deletion to have a substantial effect on GR binding, and others have demonstrated the importance of the bZIP transcription factor C/EBP (5). HNF6 (*Hnf6*), is a lineage-determining factor which is part of the onecut family of transcription factors (38). Its motif is also enriched at hepatic GR binding sites (Fig.1C), but found at a smaller proportion of GR sites than the HNF4 motif, and its presence in the vicinity of GREs is not associated with the large increase in chromatin accessibility seen with the HNF4 motif (Fig.S3A).

We therefore hypothesised that HNF6 plays a more minor role in shaping liver GR action. We used a mouse model of postnatal liver *Hnf6* deletion (its embryonic loss is lethal) (Fig.S3B). We found that this had only a small effect on the liver transcriptome under basal conditions (Fig.S3C), with a correspondingly minor impact on glucocorticoid-responsiveness. Analysis with stageR detected 148 genes with a significant treatment-genotype interaction, of which 34 showed an altered direction of significant change with glucocorticoid treatment (Fig.S3D). These data suggest that HNF6 is indeed less influential than HNF4A in shaping the response to glucocorticoid, with a lesser functional impact evident. By contrast, HNF4A is clearly critical in determining tissue-specificity of GR action. We suggest that, as a lineage-determining factor, HNF4A confers tissue-specificity to the liver GR cistrome by maintaining chromatin accessibility at HNF4 motif-marked sites (assisted loading). In the absence of HNF4A, the regulatory landscape is remodelled, and GR binds to strong canonical GRE motifs normally inaccessible in the terminally differentiated hepatocyte.

## Discussion

In this study, we show that a substantial portion of the liver GR cistrome is characterised by HNF4A binding and the HNF4 motif. The presence of the HNF4 motif favours open chromatin, in comparison to sites where the HNF4 motif is not present. Strikingly, when HNF4A is removed, the GR cistrome is remodelled, with the HNF4 motif enriched at sites where GR binding is lost. New GR binding emerges, at sites where GR is typically bound in non-liver tissues, where chromatin is not normally accessible in liver.

Multiple previous studies have demonstrated the presence of the HNF4 motif at GR binding sites (17, 21, 24, 25), and have shown the tissue-specificity of nuclear receptor cistromes (4, 18). CCAAT enhancer binding protein beta (C/EBPB) and the basic helix-loop-helix factor E47 have also been shown to play important roles in regulating hepatic glucocorticoid action (5, 25). This study builds on these works by directly comparing the GR cistrome in *Hnf4a*-intact and *Hnf4a*-null liver, identifying those GR binding sites which are dependent on HNF4A to be maintained, and showing that loss of tissue-specificity extends to the emergence of ‘non-liver’ GR binding sites. Furthermore, we show that sites marked by the HNF4 motif, and those sites lost and gained in *Hnf4a*-null liver have distinct profiles of chromatin accessibility.

The characteristics of the GR sites that are gained and lost in the absence of HNF4A suggest a balance between chromatin accessibility (4) and GRE motif strength (6) in specifying GR binding. Numbers of DNA-bound GR molecules per cell are thought to be in the orders of the hundreds (39), and are thus outnumbered by the number of potential GR binding sites (motif analysis suggesting >137,000 GREs across the mouse genome). Where HNF4A maintains chromatin accessibility, GR may bind to a weak motif with considerable degeneracy from the canonical GRE. When HNF4A is not present, GR no longer binds these sites, but can instead bind strong GRE motifs at sites where chromatin is not normally accessible in liver (Fig.S4), through its intrinsic pioneer function (6). In a similar fashion, major perturbations of the regulatory environment that likely induce chromatin remodelling (e.g. fasting (15), chronic high fat diet (17)), have been shown to alter the observed actions of GR, and we suggest that, operating through a similar mechanism, this phenomenon extends to other nuclear receptors whose activity is state-sensitive (40, 41). Indeed, ‘cistromic plasticity’ of the oestrogen receptor is proposed to be of clinical importance in breast cancer (42).

This study demonstrates, *in vivo*, the remodelling of the GR cistrome with the deletion of a lineage-determining factor. Our data echo the results of previous tissue-tissue comparisons of nuclear receptor binding (18), but now show directly the importance of a single factor. This value of the study is inextricably linked to a confounding factor, that of the abnormal liver function that results from disruption of HNF4A expression. This makes it difficult to perform more detailed physiological studies in *Hnf4a^fl/fl^Alb^Cre^* mice. However, we mitigated the liver pathology as much as possible by studying animals at a young age, in what is a widely used mouse line. The clear delineation between the lost and gained GR sites, their characteristics, and the association of these sites with glucocorticoid-responsive genes, does support an effect specific to HNF4A loss, rather than attendant liver pathology, and we do limit our conclusions to what this study tells us about GR-DNA binding.

Whilst HNF4A and GR have been identified together in ChIP-MS studies (17), our data suggest a permissive role for HNF4A akin to what is proposed for C/EBPB - that of maintaining chromatin accessibility at commonly occupied sites - rather than direct co-operative interaction between the two nuclear receptors. There is a broad distribution of intermotif distances, with many GRE-HNF4 motif pairs lying further apart than the 20bp proposed for high-confidence co-occupancy (and thus physical co-operativity) (26). There are clearly many sites where GR binding is not dependent on HNF4A, and more dynamic context-specificity of GR action will also be conferred by the ultradian and circadian variation in the availability of its endogenous ligand (43, 44). Thus, the combinatorial actions of lineage-determining factors, state-sensitive factors or chromatin remodelling enzymes, and the rhythmicity of its ligand, confer exquisite context-specificity to glucocorticoid receptor action, and must be taken into account when considering therapeutic applications.

## Author Contributions

Conceptualisation: ALH, TMP, MI, DWR. Investigation: ALH, TMP, DK. Methodology: FJG, MI. Resources: FJG, MI. Software: ALH, MI. Formal analysis: ALH, TMP, MI. Writing - original draft: ALH. Writing review & editing: ALH, MI, DAB, ASIL, DWR. Funding acquisition: ALH, ASIL, DWR. Supervision: DAB, ASIL, MI, DWR.

The authors declare no competing interests.

## ACKNOWLEDGEMENTS

We acknowledge our funders - the MRC (Clinical Research Training Fellowship MR/N021479/1 to ALH) and the Wellcome Trust (107849/Z/15/Z, 107851/Z/15/Z to ASIL and DWR). We acknowledge too the valuable contribution of the University of Manchester core facilities: the Genomic Technologies Core Facility (Andy Hayes, Stacey Holden, Claire Morrisroe), the Bioinformatics Core Facility (Rachel Scholey, Peter Briggs), the Genome Editing Unit (Maj Simonsen Jackson, Neil Humphreys), and the Biological Services Facility (Sarah Dinning, Adam Johnson). We also thank colleagues Robert Maidstone, Nicola Begley, Suzanna Dickson, and Matthew Baxter for practical assistance with experiments and data analysis.

## Methods

### Animals

Male mice were used throughout, to eliminate sex as a confounder. All animals had ad libitum access to standard laboratory chow and water, and were group-housed on 12hr:12hr light-dark cycles. All experiments on wild-type and *Hnf6^fl/fl^Alb^CreERT2+/−^* mice were conducted at the University of Manchester in accordance with local requirements and with the UK Animals (Scientific Procedures) Act 1986. Procedures were approved by the University of Manchester Animal Welfare and Ethical Review Body (AWERB) and carried out under licence (project licence 70/8558, held by DAB). Dexamethasone and vehicle treatment of *Hnf4a^fl/fl^Alb^Cre+/−^* mice was carried out at the National Cancer Institute as described in (21). The National Cancer Institute Animal Care and Use Committee approved all animal experiments conducted in these experiments.

Wild-type C57BL/6 mice (Figure 1) were obtained from an in-house colony. *Hnf6^fl/fl^Alb^CreERT2+/−^* mi ce were generated in-house using the *Onecut1^tm1.1Mga^*/Mmnc (*Hnf6^fl/fl^*) line (45) (sperm obtained from MMRRC) and the *Alb^tm1(cre/ERT2)Mtz^* line (46) (kindly gifted by Drs Pierre Chambon and Daniel Metzger. Recombination was induced with tamoxifen as described (41).

### Glucocorticoid administration

For acute treatment with dexamethasone, mice were injected by the intraperitoneal route with water-soluble dexamethasone (D2915 - Sigma-Aldrich) at a dose of 1mg/kg, dissolved in water for injection to a final dexamethasone concentration of 0.2mg/ml. Corresponding vehicle treatment was an equivalent mass of (2-hydroxypropyl)-*β*-cyclodextrin (H107 - Sigma-Aldrich) dissolved in water for injection. For studies of GR binding or chromatin accessibility (ChIP-seq and ATAC-seq), tissue was collected after one or two hours; for studies of gene expression (RNA-seq), tissue was collected after two hours.

### Chromatin immunoprecipitation (ChIP)

#### Tissue processing and chromatin preparation

Chromatin was prepared from flash-frozen liver tissue using the Active Motif ChIP-IT High Sensitivity kit (Active Motif), employing a modified protocol described in (47). All ChIP experiments were conducted with two biological replicates per group, as per ENCODE standards, with replicates handled separately through *in vivo* to *in silico* steps.

#### Immunoprecipitation (IP) and DNA elution

25*μ*g of chromatin (using Nanodrop-measured concentration) was incubated overnight at 4°Cwith a GR antibody cocktail (Protein-Tech 24050-1-A (lot 00044414) and Cell Signaling D8H2 (lot 2) (2*μ*l of each per IP reaction)). As described (47), to permit direct comparison between samples, and to control for technical variation between ChIP reactions, a spike-in ChIP normalisation strategy was employed (31), with 30ng spike-in chromatin (53083, Active Motif) and 2*μ*g spike-in antibody (61686, Active Motif (lot 34216004)) being included in each IP reaction. To obtain sufficient DNA for next-generation sequencing, three IP reactions were carried out for each sample, and then pooled for the pull-down step. Antibody was pulled down using 10*μ*l washed magnetic protein G agarose beads (ReSyn Biosciences). Beads were washed five times with AM1 Wash Buffer (Active Motif) then DNA eluted as per ChIP-IT kit instructions. ChIP-seq DNA was purified with the MinElute PCR Purification kit (Qiagen) (two 10*μ*l elutions per sample).

#### Library preparation

Library preparation and sequencing steps were carried out by the University of Manchester Genomic Technologies Core Facility, using the TruSeq® ChIP library preparation kit (Illumina) and subsequent paired-end sequencing on the Illumina HiSeq 4000 platform.

#### Raw data processing

Raw FASTQ files were quality checked with FastQC software (v0.11.7, [Babraham Bioinformatics](http://www.bioinformatics.babraham.ac.uk/projects/fastqc/)). Reads were then trimmed with Trimmomatic (v0.38) and aligned to the reference genomes (mouse (mm10) and drosophila (dm6) as appropriate) with Bowtie2 (v2.3.4.3) (49). The resulting SAM (Sequence Alignment Map) files were converted to BAM (Binary Alignment Map) files, sorted and indexed with SAMtools (v1.9) (50). Duplicates were removed with Picard (v2.18.14, [Broad Institute](https://broadinstitute.github.io/picard/)). For published ChIP-seq and DNase-seq data, the sratoolkit package (v2.9.2, [NCBI](https://trace.ncbi.nlm.nih.gov/Traces/sra/sra.cgi?view=software)) was used to download FASTQ files from the GEO Sequence Read Archive. These were then processed as above.

### Assay for transposase-accessible chromatin (ATAC)

ATAC-seq service was performed on frozen liver tissue by Active Motif, on two biological replicates. Libraries were sequenced (paired end) on the Illumina NextSeq 500 platform. Reads were aligned to mm10, and SAMtools used to create sorted, indexed BAM files as above.

### RNA sequencing (RNA-seq)

#### Sample and library preparation

RNA extraction from liver tissue (n=3-6 biological replicates per group) was performed using the ReliaPrep RNA Miniprep system (Promega), as per manufacturer’s instructions, incorporating a DNase treatment step. Lysing Matrix D tubes (MP Biomedicals) were used to homogenise tissue. 1*μ*g RNA was supplied to the Genomic Technologies Core Facility for library preparation and paired-end sequencing on the Illumina HiSeq 4000 platform, with the TruSeq® Stranded mRNA assay kit (Illumina) employed as per manufacturer’s instructions. Demultiplexing (one mismatch allowed) and BCL-to-Fastq conversion was carried out using bcl2fastq software (v2.17.1.14) (Illumina).

#### Raw data processing

FASTQ files were processed by the Core Bioinformatics Facility, employing FastQC Screen (v0.9.2) (51). Trimmomatic (v0.36) (48) was used to remove adapters and poor quality bases. STAR (v2.5.3a) (52) was used to map reads to the mm10 reference genome; counts per gene (exons, GENCODEM16) were then used in differential expression analysis (see below).

### qPCR

RNA was converted to cDNA with the High Capacity RNA-to-cDNA kit (Applied Biosystems). qPCR was performed with PowerUp SYBR Green Master Mix (Thermo Fisher Scientific) using the StepOne Plus (Applied Biosystems) platform. Expression of *Hnf6* (forward primer: GGCAACGTGAGCGGTAGTTT; reverse primer: TTGCTGGGAGTTGTGAATGCT) and *Hnf4a* (forward: AGAAGATTGCCAACATCAC; reverse: GGTCATCCA-GAAGGAGTT) was normalised to *Actb* (forward: GGCTG-TATTCCCCTCCATCG; reverse: CCAGTTGGTAACAAT-GCCATGT).

### Data analysis

#### ChIP-seq peak-calling

Peak-calling was performed using MACS2 (v2.1.1.20160309) (53) with settings for narrow peaks and with a q-value cut-off of 0.01 (parameters set as: -f BAMPE -g mm –keep-dup=1 -q 0.01 –bdg –SPMR –verbose 0).

#### ChIP-seq differential binding (DB) analysis

This was performed with *csaw* (v1.20.0) (29, 30), incorporating spike-in normalisation, as per the code uploaded to [Mendeley Data](https://data.mendeley.com/datasets/k8d386ndz6/2).

#### ChIP-seq and ATAC-seq coverage

deepTools (bamCoverage and computeMatrix commands) (54) was used to determine read coverage over regions of interest. The computeMatrix command was used in reference-point mode, with the reference point set as the centre of each region.

#### Peak annotation and motif analysis

HOMER (v4.9.1) (55) was used to annotate peak locations (annotatePeaks.pl). HOMER was also used to analyse the underlying DNA sequence motifs in either MACS2-called peaks or *csaw*-defined DB sites. The findMotifsGenome.pl package was used for motif enrichment analysis, with window size set to 200bp (default), and the -mask option set. To determine abundance of specific motifs within a set of regions, we used annotatePeaks.pl with the -m and -hist options set; to determine the scores of GRE motifs detected, we used findMotifsGenome.pl with the -find option. To detect instances of motifs genome-wide, we used HOMER’s scanMotifGenomeWide.pl package. Throughout, we used the “GRE(NR),IR3/A549-GR-ChIP-Seq(GSE32465)/Homer” matrix as representative of the canonical GRE, the “HNF4a(NR),DR1/HepG2-HNF4a-ChIP-Seq(GSE25021)/Homer” matrix for the HNF4 motif, and “HNF6(Homeobox)/Liver-Hnf6-ChIP-Seq(ERP000394)/Homer” for HNF6; motif matrices are available at http://homer.ucsd.edu/homer/custom.motifs. HOMER motif files specify a log odds detection threshold; this was left unaltered for detection of ‘strong’ motifs, and reduced by 3 for detection of ‘weak’ motifs.

#### Distances between peaks (or motifs)

bedtools (56) (intersect and window tools) was used to determine overlap between peak sets, or to determine distances between peaks or motifs.

#### Overlap with CistromeDB database

GIGGLE (35) (acccessed through the CistromeDB Toolkit portal) was used to look for overlap of sites of interest with published datasets (top 1k peaks in each dataset). The tool was set to apply to the mm10 genome, and transcription factor data.

#### Visualisation of ChIP-seq data

Visualisations of ChIP-seq signal tracks were created with Integrative Genomics Viewer (IGV) (57) and deepTools (54).

#### Differential gene expression and pathway analysis

Differentially expressed genes were identified with edgeR (v3.28.1) (58, 59), and detection of a genotype-treatment interaction effect was performed with limma (v3.42.2) voom (60) and stageR (v1.8.0) (37), as per the code uploaded to [Mendeley Data](https://data.mendeley.com/datasets/k8d386ndz6/2). Pathway enrichment analyses were performed with ReactomePA (61), using *enrichPathway(genes, organism = “mouse”, pvalueCutoff = 0.05, pAdjustMethod = “BH”, qvalueCutoff = 0.1, maxGSSize = 2000, readable = FALSE)*.

#### Integration of ChIP-seq and RNA-seq data

PEGS (Peak-set Enrichment of Gene-Sets) (https://github.com/fls-bioinformatics-core/pegs) was employed to calculate enrichment (hypergeometric test) of genes of interest within specified distances of peak sets. The genome was set to mm10, and distances (bp) specified as 100, 500, 1000, 5000, 10000, 50000, 100000, 500000, 1000000, 5000000.

### Published datasets used

The following datasets were downloaded from the GEO Sequence Read Archive: ZT6 liver DNase-seq (SRR1551954) (32), mouse liver H3K27ac ChIP-seq (SRR5054771) (33), mouse liver H3K4me1 ChIP-seq (SRR317236, SRR317235) (34), mouse liver H3K27me3 ChIP-seq (SRR566941, SRR566942) (34), mouse liver HNF4A ChIP-seq (SRR7634103, SRR7634104, SRR7634105, SRR3151870, SRR3151871, SRR3151878, SRR3151879) (62, 63), and mouse macrophage GR ChIP-seq (SRR5182692) (11).

### Plots and statistics

Plots were created with ggplot2, incorporating statistical tests by ggpubr, or with GraphPad Prism v8.

## Data and Code Availability

Sequencing data is available through [ArrayExpress](https://www.ebi.ac.uk/arrayexpress/) at the following accession numbers: ChIP-seq - E-MTAB-10224; RNA-seq - E-MTAB-10247; ATAC-seq - E-MTAB-10266. Outputs of peak-calling, differential binding analysis, differential expression analyses, HOMER motif discovery and GIGGLE analyses have been uploaded to Mendeley Data doi:10.17632/k8d386ndz6.2, as has the R code for differential binding and differential expression analyses. The PEGS Python package is freely available at https://github.com/fls-bioinformatics-core/pegs.

## SUPPLEMENTARY INFORMATION: Supplementary Figures

**Fig. S1.**
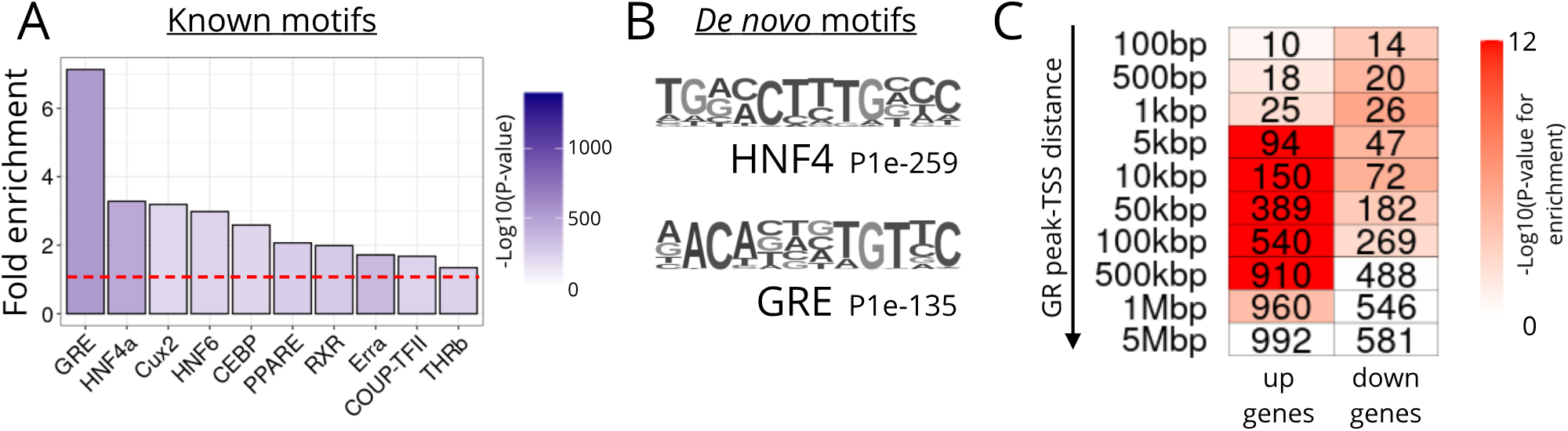
**A.** Fold enrichment, in GR ChIP-seq peaks from vehicle-treated mouse liver, of known motifs. Red dotted line at y=1. **B.** The two motifs detected most strongly (lowest P values) *de novo* in GR peaks. **C.** Heatmap showing enrichment (hypergeometric test) of the transcription start sites (TSSs) of genes up or downregulated by glucocorticoid treatment at increasing distances from GR ChIP-seq peaks (VEH samples). Shading of each cell indicates −log10(P-value) for enrichment (over all genes in the genome), number indicates number of genes in each cluster at that distance.

**Fig. S2.**
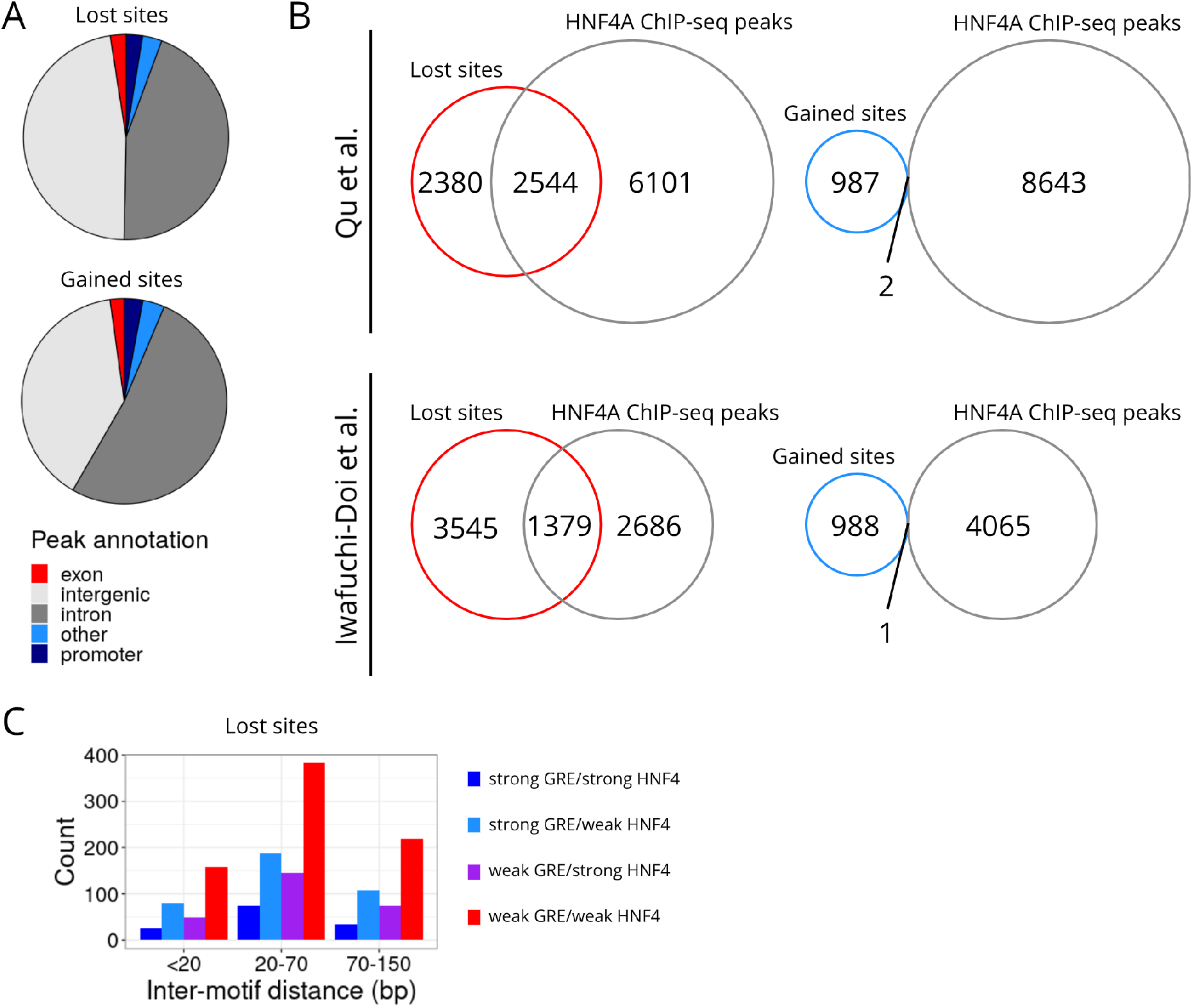
**A.** Piecharts showing annotated locations of GR sites lost (top) and gained (bottom) with *Hnf4a* deletion. **B.** Venn diagrams showing overlap of lost and gained GR sites with published HNF4A cistromes from (62) (top) and (63) (bottom). **C.** Barchart of intermotif distances for GRE and HNF4 motifs detected within lost GR sites.

**Fig. S3.**
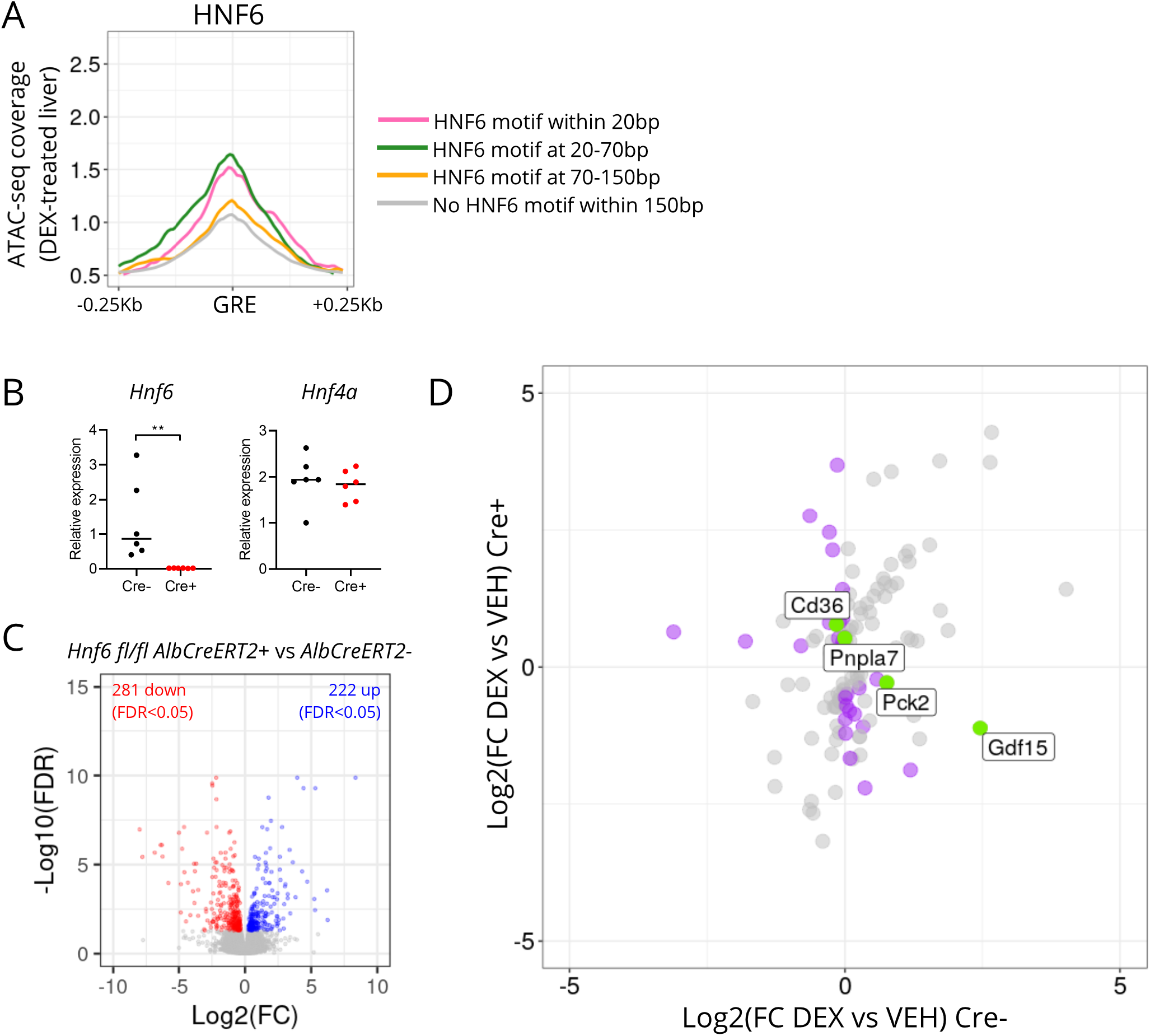
**A.** ATAC-seq coverage score, in DEX-treated liver, around canonical GRE motifs with or without a HNF6 motif (left panel), or HNF4 motif (right panel, duplicate of Figure 1F, provided here for comparison), within specified distances. **B.** Liver expression of *Hnf6* and *Hnf4a* (as determined by qPCR) in *Hnf6^fl/fl^Alb^CreERT2+/−^* mice. n=6 per group, line at median. **P<0.01, Mann Whitney test. **C.** Liver RNA-seq in *Hnf6^fl/fl^Alb^CreERT2^* mice, vehicle-treated Cre+ vs vehicle-treated Cre- samples. Significantly downregulated genes (FDR<0.05) in red, significantly upregulated genes in blue. **D.** Effect of DEX treatment in Cre- and Cre+ mice. Genes where stageR detects a significant treatment x genotype interaction shown. Those where direction of (significant) change is different between genotypes highlighted in purple. These include metabolic regulators and enzymes of interest, highlighted in green.

**Fig. S4.**
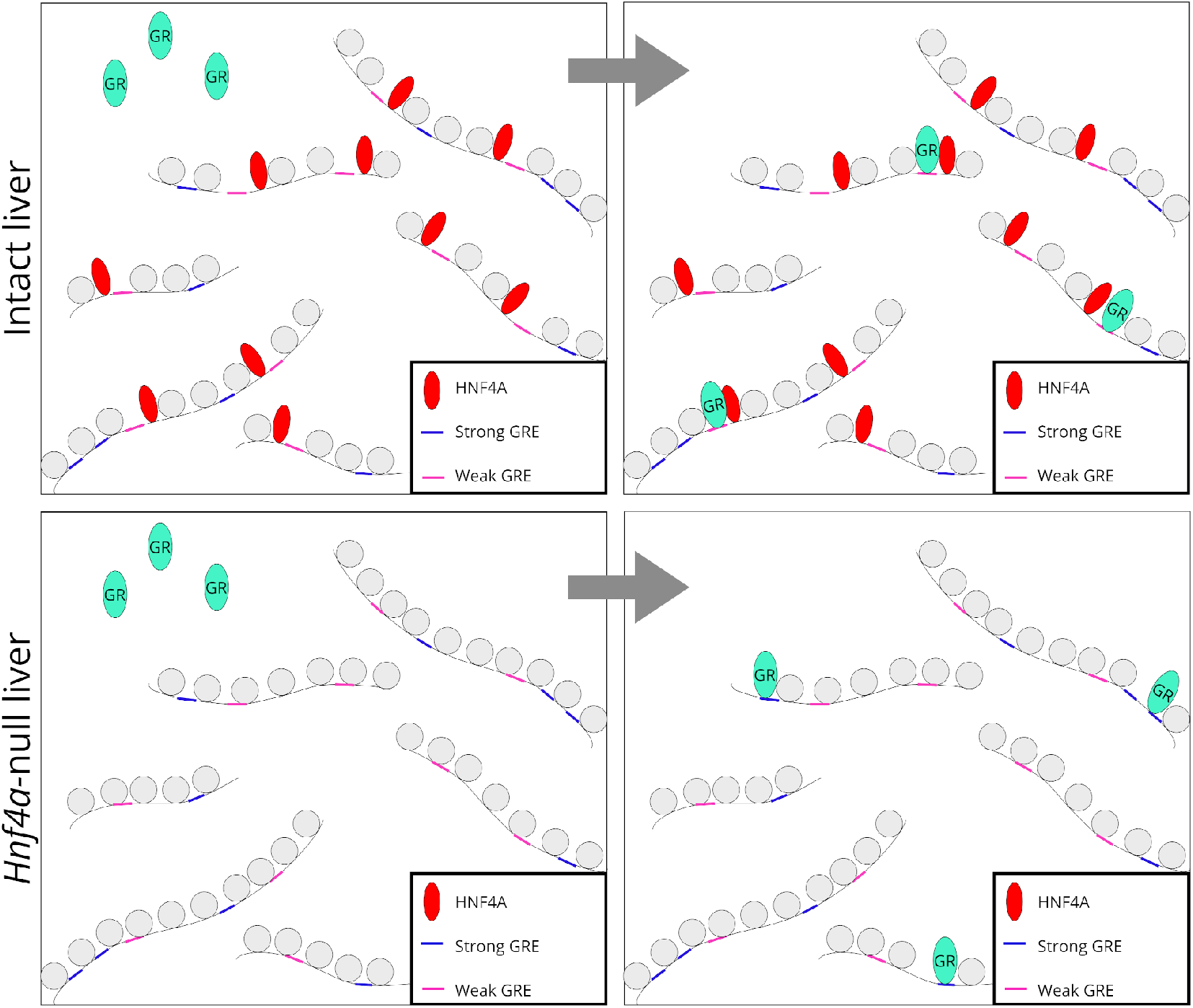
Cartoon. Proposed patterns of GR binding in intact (*Hnf4a^fl/fl^Alb^Cre^* Cre-) and *Hnf4a*-null (*Hnf4a^fl/fl^Alb^Cre^* Cre-) mouse liver in the course of glucocorticoid treatment. In intact liver, HNF4A binding marks sites where open chromatin favours GR binding, even though GREs may show considerable degeneracy from the canonical motif (“Weak GREs“). In *Hnf4a*-null liver, greater similarity to the canonical GRE (“Strong GRES“) favours GR binding, as HNF4A-mediated chromatin accessibility is lost.

## References

1. Mary F Dallman. Fast glucocorticoid actions on brain: back to the future. Front. Neuroendocrinol., 26(3–4):103–108, October 2005.

2. Stephen Kershaw, David J Morgan, James Boyd, David G Spiller, Gareth Kitchen, Egor Zindy, Mudassar Iqbal, Magnus Rattray, Christopher M Sanderson, Andrew Brass, Claus Jorgensen, Tracy Hussell, Laura C Matthews, and David W Ray. Glucocorticoids rapidly inhibit cell migration through a novel, non-transcriptional HDAC6 pathway. J. Cell Sci., 133 (11), June 2020.

3. Thomas A Johnson, Ville Paakinaho, Sohyoung Kim, Gordon L Hager, and Diego M Presman. Genome-wide binding potential and regulatory activity of the glucocorticoid receptor’s monomeric and dimeric forms. Nat. Commun., 12(1), December 2021.

4. Sam John, Peter J Sabo, Robert E Thurman, Myong-Hee Sung, Simon C Biddie, Thomas A Johnson, Gordon L Hager, and John A Stamatoyannopoulos. Chromatin accessibility pre-determines glucocorticoid receptor binding patterns. Nat. Genet., 43(3):264–268, March 2011.

5. Lars Grøntved, Sam John, Songjoon Baek, Ying Liu, John R Buckley, Charles Vinson, Greti Aguilera, and Gordon L Hager. C/EBP maintains chromatin accessibility in liver and facilitates glucocorticoid receptor recruitment to steroid response elements. EMBO J., 32 (11):1568–1583, May 2013.

6. Thomas A Johnson, Razvan V Chereji, Diana A Stavreva, Stephanie A Morris, Gordon L Hager, and David J Clark. Conventional and pioneer modes of glucocorticoid receptor interaction with enhancer chromatin in vivo. Nucleic Acids Res., 46(1):203–214, January 2018.

7. Ian C McDowell, Alejandro Barrera, Anthony M D’Ippolito, Christopher M Vockley, Linda K Hong, Sarah M Leichter, Luke C Bartelt, William H Majoros, Lingyun Song, Alexias Safi, D Dewran Koçak, Charles A Gersbach, Alexander J Hartemink, Gregory E Crawford, Barbara E Engelhardt, and Timothy E Reddy. Glucocorticoid receptor recruits to enhancers and drives activation by motif-directed binding. Genome Res., 28(9):1272–1284, September 2018.

8. Anthony M D’Ippolito, Ian C McDowell, Alejandro Barrera, Linda K Hong, Sarah M Leichter, Luke C Bartelt, Christopher M Vockley, William H Majoros, Alexias Safi, Lingyun Song, Charles A Gersbach, Gregory E Crawford, and Timothy E Reddy. Pre-established chromatin interactions mediate the genomic response to glucocorticoids. Cell Syst, 7(2):146–160.e7, August 2018.

9. Tina B Miranda, Ty C Voss, and Gordon L Hager. High-throughput fluorescence-based screen to identify factors involved in nuclear receptor recruitment to response elements. Methods Mol. Biol., 1042:3–12, 2013.

10. Maria A Sacta, Bowranigan Tharmalingam, Maddalena Coppo, David A Rollins, Dinesh K Deochand, Bradley Benjamin, Li Yu, Bin Zhang, Xiaoyu Hu, Rong Li, Yurii Chinenov, and Inez Rogatsky. Gene-specific mechanisms direct glucocorticoid-receptor-driven repression of inflammatory response genes in macrophages. Elife, 7, February 2018.

11. Kyu-Seon Oh, Heta Patel, Rachel A Gottschalk, Wai Shing Lee, Songjoon Baek, Iain D C Fraser, Gordon L Hager, and Myong-Hee Sung. Anti-Inflammatory chromatinscape suggests alternative mechanisms of glucocorticoid receptor action. Immunity, 47(2):298–309.e5, August 2017.

12. Laura Escoter-Torres, Franziska Greulich, Fabiana Quagliarini, Michael Wierer, and Nina Henriette Uhlenhaut. Anti-inflammatory functions of the glucocorticoid receptor require DNA binding. Nucleic Acids Res., 48(15):8393–8407, September 2020.

13. Milan Surjit, Krishna Priya Ganti, Atish Mukherji, Tao Ye, Guoqiang Hua, Daniel Metzger, Mei Li, and Pierre Chambon. Widespread negative response elements mediate direct repression by agonist-liganded glucocorticoid receptor. Cell, 145(2):224–241, April 2011.

14. William H Hudson, Ian Mitchelle S de Vera, Jerome C Nwachukwu, Emily R Weikum, Austin G Herbst, Qin Yang, David L Bain, Kendall W Nettles, Douglas J Kojetin, and Eric A Ortlund. ryptic glucocorticoid receptor-binding sites pervade genomic NF-κB response elements. Nat. Commun., 9(1):1337, April 2018.

15. Ido Goldstein, Songjoon Baek, Diego M Presman, Ville Paakinaho, Erin E Swinstead, and Gordon L Hager. Transcription factor assisted loading and enhancer dynamics dictate the hepatic fasting response. Genome Res., 27(3):427–439, March 2017.

16. N Henriette Uhlenhaut, Grant D Barish, Ruth T Yu, Michael Downes, Malith Karunasiri, Christopher Liddle, Petra Schwalie, Norbert Hübner, and Ronald M Evans. Insights into negative regulation by the glucocorticoid receptor from genome-wide profiling of inflammatory cistromes. Mol. Cell, 49(1):158–171, January 2013.

17. Fabiana Quagliarini, Ashfaq Ali Mir, Kinga Balazs, Michael Wierer, Kenneth Allen Dyar, Celine Jouffe, Konstantinos Makris, Johann Hawe, Matthias Heinig, Fabian Volker Filipp, Grant Daniel Barish, and Nina Henriette Uhlenhaut. Cistromic reprogramming of the diurnal glucocorticoid hormone response by High-Fat diet. Mol. Cell, 76(4):531–545.e5, November 2019.

18. Jason Gertz, Daniel Savic, Katherine E Varley, E Christopher Partridge, Alexias Safi, Preti Jain, Gregory M Cooper, Timothy E Reddy, Gregory E Crawford, and Richard M Myers. Distinct properties of cell-type-specific and shared transcription factor binding sites. Mol. Cell, 52(1):25–36, October 2013.

19. Joshua R Beytebiere, Alexandra J Trott, Ben J Greenwell, Collin A Osborne, Helene Vitet, Jessica Spence, Seung-Hee Yoo, Zheng Chen, Joseph S Takahashi, Noushin Ghaffari, and Jerome S Menet. Tissue-specific BMAL1 cistromes reveal that rhythmic transcription is associated with rhythmic enhancer-enhancer interactions. Genes Dev., 33(5–6):294–309, March 2019.

20. Michael I Love, Matthew R Huska, Marcel Jurk, Robert Schöpflin, Stephan R Starick, Kevin Schwahn, Samantha B Cooper, Keith R Yamamoto, Morgane Thomas-Chollier, Martin Vingron, and Sebastiaan H Meijsing. Role of the chromatin landscape and sequence in determining cell type-specific genomic glucocorticoid receptor binding and gene regulation. Nucleic Acids Res., 45(4):1805–1819, February 2017.

21. Giorgio Caratti, Mudassar Iqbal, Louise Hunter, Donghwan Kim, Ping Wang, Ryan M Vonslow, Nicola Begley, Abigail J Tetley, Joanna L Woodburn, Marie Pariollaud, Robert Maidstone, Ian J Donaldson, Zhenguang Zhang, Louise M Ince, Gareth Kitchen, Matthew Baxter, Toryn M Poolman, Dion A Daniels, David R Stirling, Chad Brocker, Frank Gonzalez, Andrew Si Loudon, David A Bechtold, Magnus Rattray, Laura C Matthews, and David W Ray. REVERBa couples the circadian clock to hepatic glucocorticoid action. J. Clin. Invest., 128(10):4454–4471, October 2018.

22. Shen-Hsi Yang, Munazah Andrabi, Rebecca Biss, Syed Murtuza Baker, Mudassar Iqbal, and Andrew D Sharrocks. ZIC3 controls the transition from naive to primed pluripotency. Cell Rep., 27(11):3215–3227.e6, June 2019.

23. Alasdair W Jubb, Robert S Young, David A Hume, and Wendy A Bickmore. Enhancer turnover is associated with a divergent transcriptional response to glucocorticoid in mouse and human macrophages. J. Immunol., 196(2):813–822, January 2016.

24. Hee-Woong Lim, N Henriette Uhlenhaut, Alexander Rauch, Juliane Weiner, Sabine Hübner, Norbert Hübner, Kyoung-Jae Won, Mitchell A Lazar, Jan Tuckermann, and David J Steger. Genomic redistribution of GR monomers and dimers mediates transcriptional response to exogenous glucocorticoid in vivo. Genome Res., 25(6):836–844, June 2015.

25. M Charlotte Hemmer, Michael Wierer, Kristina Schachtrup, Michael Downes, Norbert Hübner, Ronald M Evans, and N Henriette Uhlenhaut. E47 modulates hepatic glucocorticoid action. Nat. Commun., 10(1):306, January 2019.

26. Can Sönmezer, Rozemarijn Kleinendorst, Dilek Imanci, Guido Barzaghi, Laura Villacorta, Dirk Schübeler, Vladimir Benes, Nacho Molina, and Arnaud Regis Krebs. Molecular co-occupancy identifies transcription factor binding cooperativity in vivo. Mol. Cell, 81(2):255–267.e6, January 2021.

27. Stine M Præstholm, Catarina M Correia, and Lars Grøntved. Multifaceted control of GR signaling and its impact on hepatic transcriptional networks and metabolism. Front. Endocrinol., 11:572981, October 2020.

28. Graham P Hayhurst, Ying-Hue Lee, Gilles Lambert, Jerrold M Ward, and Frank J Gonzalez. Hepatocyte nuclear factor 4alpha (nuclear receptor 2a1) is essential for maintenance of hepatic gene expression and lipid homeostasis. Mol. Cell. Biol., 21(4):1393–1403, February 2001.

29. Aaron T L Lun and Gordon K Smyth. csaw: a bioconductor package for differential binding analysis of ChIP-seq data using sliding windows. Nucleic Acids Res., 44(5):e45, March 2016.

30. Aaron T L Lun and Gordon K Smyth. From reads to regions: a bioconductor workflow to detect differential binding in ChIP-seq data. F1000Res., 4:1080, October 2015.

31. Brian Egan, Chih-Chi Yuan, Madeleine Lisa Craske, Paul Labhart, Gulfem D Guler, David Arnott, Tobias M Maile, Jennifer Busby, Chisato Henry, Theresa K Kelly, Charles A Tindell, Suchit Jhunjhunwala, Feng Zhao, Charlie Hatton, Barbara M Bryant, Marie Classon, and Patrick Trojer. An alternative approach to ChIP-Seq normalization enables detection of Genome-Wide changes in histone H3 lysine 27 trimethylation upon EZH2 inhibition. PLoS One, 11(11):e0166438, November 2016.

32. Jonathan Aryeh Sobel, Irina Krier, Teemu Andersin, Sunil Raghav, Donatella Canella, Federica Gilardi, Alexandra Styliani Kalantzi, Guillaume Rey, Benjamin Weger, Frédéric Gachon, Matteo Dal Peraro, Nouria Hernandez, Ueli Schibler, Bart Deplancke, Felix Naef, and CycliX consortium. Transcriptional regulatory logic of the diurnal cycle in the mouse liver. PLoS Biol., 15(4):e2001069, April 2017.

33. Sean M Armour, Jarrett R Remsberg, Manashree Damle, Simone Sidoli, Wesley Y Ho, Zhenghui Li, Benjamin A Garcia, and Mitchell A Lazar. An HDAC3-PROX1 corepressor module acts on HNF4α to control hepatic triglycerides. Nat. Commun., 8(1):549, September 2017.

34. Yin Shen, Feng Yue, David F McCleary, Zhen Ye, Lee Edsall, Samantha Kuan, Ulrich Wagner, Jesse Dixon, Leonard Lee, Victor V Lobanenkov, and Bing Ren. A map of the cisregulatory sequences in the mouse genome. Nature, 488(7409):116–120, August 2012.

35. Ryan M Layer, Brent S Pedersen, Tonya DiSera, Gabor T Marth, Jason Gertz, and Aaron R Quinlan. GIGGLE: a search engine for large-scale integrated genome analysis. Nat. Methods, 15(2):123–126, February 2018.

36. Yong Hoon Kim, Sajid A Marhon, Yuxiang Zhang, David J Steger, Kyoung-Jae Won, and Mitchell A Lazar. Rev-erbα dynamically modulates chromatin looping to control circadian gene transcription. Science, 359(6381):1274–1277, March 2018.

37. Koen Van den Berge, Charlotte Soneson, Mark D Robinson, and Lieven Clement. stager: a general stage-wise method for controlling the gene-level false discovery rate in differential expression and differential transcript usage. Genome Biol., 18(1):151, August 2017.

38. Duncan T Odom, Nora Zizlsperger, D Benjamin Gordon, George W Bell, Nicola J Rinaldi, Heather L Murray, Tom L Volkert, Jörg Schreiber, P Alexander Rolfe, David K Gifford, Ernest Fraenkel, Graeme I Bell, and Richard A Young. Control of pancreas and liver gene expression by HNF transcription factors. Science, 303(5662):1378–1381, February 2004.

39. Ville Paakinaho, Diego M Presman, David A Ball, Thomas A Johnson, R Louis Schiltz, Peter Levitt, Davide Mazza, Tatsuya Morisaki, Tatiana S Karpova, and Gordon L Hager. Single-molecule analysis of steroid receptor and cofactor action in living cells. Nat. Commun., 8: 15896, June 2017.

40. Yuxiang Zhang, Romeo Papazyan, Manashree Damle, Bin Fang, Jennifer Jager, Dan Feng, Lindsey C Peed, Dongyin Guan, Zheng Sun, and Mitchell A Lazar. The hepatic circadian clock fine-tunes the lipogenic response to feeding through RORα/γ. Genes Dev., 31(12): 1202–1211, June 2017.

41. A Louise Hunter, Charlotte E Pelekanou, Antony Adamson, Polly Downton, Nichola J Barron, Thomas Cornfield, Toryn M Poolman, Neil Humphreys, Peter S Cunningham, Leanne Hodson, Andrew S I Loudon, Mudassar Iqbal, David A Bechtold, and David W Ray. Nuclear receptor REVERB α is a state-dependent regulator of liver energy metabolism. Proc. Natl. Acad. Sci. U. S. A., 117(41):25869–25879, October 2020.

42. Isabel Mayayo-Peralta, Stefan Prekovic, and Wilbert Zwart. Estrogen receptor on the move: Cistromic plasticity and its implications in breast cancer. Mol. Aspects Med., (100939): 100939, December 2020.

43. Becky L Conway-Campbell, John R Pooley, Gordon L Hager, and Stafford L Lightman. Molecular dynamics of ultradian glucocorticoid receptor action. Mol. Cell. Endocrinol., 348 (2):383–393, January 2012.

44. Louise M Ince, Zhenguang Zhang, Stephen Beesley, Ryan M Vonslow, Ben R Saer, Laura C Matthews, Nicola Begley, Julie E Gibbs, David W Ray, and Andrew S I Loudon. Circadian variation in pulmonary inflammatory responses is independent of rhythmic glucocorticoid signaling in airway epithelial cells. FASEB J., 33(1):126–139, January 2019.

45. Hongjie Zhang, Elizabeth Tweedie Ables, Christine F Pope, M Kay Washington, Susan Hipkens, Anna L Means, Gunter Path, Jochen Seufert, Robert H Costa, Andrew B Leiter, Mark A Magnuson, and Maureen Gannon. Multiple, temporal-specific roles for HNF6 in pancreatic endocrine and ductal differentiation. Mech. Dev., 126(11-12):958–973, December 2009.

46. Michael Schuler, Andrée Dierich, Pierre Chambon, and Daniel Metzger. Efficient temporally controlled targeted somatic mutagenesis in hepatocytes of the mouse. Genesis, 39(3): 167–172, 2004.

47. Ann Louise Hunter, Natasha Narang, Matthew Baxter, David W Ray, and Toryn M Poolman. An improved method for quantitative ChIP studies of nuclear receptor function. J. Mol. Endocrinol., 62(4):169–177, May 2019.

48. Anthony M Bolger, Marc Lohse, and Bjoern Usadel. Trimmomatic: a flexible trimmer for illumina sequence data. Bioinformatics, 30(15):2114–2120, August 2014.

49. Ben Langmead and Steven L Salzberg. Fast gapped-read alignment with bowtie 2. Nat. Methods, 9(4):357–359, March 2012.

50. Heng Li, Bob Handsaker, Alec Wysoker, Tim Fennell, Jue Ruan, Nils Homer, Gabor Marth, Goncalo Abecasis, Richard Durbin, and 1000 Genome Project Data Processing Subgroup. The sequence Alignment/Map format and SAMtools. Bioinformatics, 25(16):2078–2079, August 2009.

51. Steven W Wingett and Simon Andrews. FastQ screen: A tool for multi-genome mapping and quality control. F1000Res., 7:1338, August 2018.

52. Alexander Dobin, Carrie A Davis, Felix Schlesinger, Jorg Drenkow, Chris Zaleski, Sonali Jha, Philippe Batut, Mark Chaisson, and Thomas R Gingeras. STAR: ultrafast universal RNA-seq aligner. Bioinformatics, 29(1):15–21, January 2013.

53. Yong Zhang, Tao Liu, Clifford A Meyer, Jérôme Eeckhoute, David S Johnson, Bradley E Bernstein, Chad Nusbaum, Richard M Myers, Myles Brown, Wei Li, and X Shirley Liu. Model-based analysis of ChIP-Seq (MACS). Genome Biol., 9(9):R137, September 2008.

54. Fidel Ramírez, Friederike Dündar, Sarah Diehl, Björn A Grüning, and Thomas Manke. deep-tools: a flexible platform for exploring deep-sequencing data. Nucleic Acids Res., 42(Web Server issue):W187–91, July 2014.

55. Sven Heinz, Christopher Benner, Nathanael Spann, Eric Bertolino, Yin C Lin, Peter Laslo, Jason X Cheng, Cornelis Murre, Harinder Singh, and Christopher K Glass. Simple combinations of lineage-determining transcription factors prime cis-regulatory elements required for macrophage and B cell identities. Mol. Cell, 38(4):576–589, May 2010.

56. Aaron R Quinlan. BEDTools: The Swiss-Army tool for genome feature analysis. Curr. Protoc. Bioinformatics, 47(1):11.12.1–34, September 2014.

57. James T Robinson, Helga Thorvaldsdóttir, Wendy Winckler, Mitchell Guttman, Eric S Lander, Gad Getz, and Jill P Mesirov. Integrative genomics viewer. Nat. Biotechnol., 29(1): 24–26, January 2011.

58. Yunshun Chen, Aaron T L Lun, and Gordon K Smyth. From reads to genes to pathways: differential expression analysis of RNA-Seq experiments using rsubread and the edger quasilikelihood pipeline. F1000Res., 5:1438, June 2016.

59. Mark D Robinson, Davis J McCarthy, and Gordon K Smyth. edger: a bioconductor package for differential expression analysis of digital gene expression data. Bioinformatics, 26(1): 139–140, January 2010.

60. Charity W Law, Yunshun Chen, Wei Shi, and Gordon K Smyth. voom: Precision weights unlock linear model analysis tools for RNA-seq read counts. Genome Biol., 15(2):R29, February 2014.

61. Guangchuang Yu and Qing-Yu He. ReactomePA: an R/Bioconductor package for reactome pathway analysis and visualization. Mol. Biosyst., 12(2):477–479, February 2016.

62. Meng Qu, Tomas Duffy, Tsuyoshi Hirota, and Steve A Kay. Nuclear receptor HNF4A transrepresses CLOCK:BMAL1 and modulates tissue-specific circadian networks. Proc. Natl. Acad. Sci. U. S. A., 115(52):E12305–E12312, December 2018.

63. Makiko Iwafuchi-Doi, Greg Donahue, Akshay Kakumanu, Jason A Watts, Shaun Mahony, B Franklin Pugh, Dolim Lee, Klaus H Kaestner, and Kenneth S Zaret. The pioneer transcription factor FoxA maintains an accessible nucleosome configuration at enhancers for Tissue-Specific gene activation. Mol. Cell, 62(1):79–91, April 2016.

